# Sensitized infants who tolerate peanut have more dendritic cells and HLA-DR+ memory CD4 T cells

**DOI:** 10.64898/2026.10.20.760474

**Authors:** Charvi Cherukuri, Khodor I. Abou-Daya

## Abstract

Some infants with peanut-specific IgE tolerate peanut, but the cellular features that distinguish this sensitized tolerance from peanut allergy are poorly defined. Here we reanalyze mass cytometry data from unstimulated peripheral blood mononuclear cells of 36 one-year-old infants with challenge-confirmed peanut allergy, sensitized tolerance or neither. Clustering all 3,560,317 cells into 19 curated cell types, we find that sensitized-tolerant infants have more HLA-DR^+^ memory CD4 T cells (1.97-fold, *q* = 0.033) and lineage-negative CD38^+^ cells (2.26-fold, *q* = 0.006) than allergic infants, and a trend toward more conventional dendritic cells (2.55-fold, *q* = 0.069). The dendritic-cell difference persists in samples from the bright-staining batch, whereas the T-cell difference weakens. Of the published signatures, the expansion of plasmacytoid dendritic cells in sensitized-tolerant infants is reproduced, whereas the naive CD4 T-cell and B-cell differences track a staining batch that includes half of the sensitized-tolerant samples, in the published values as in ours. Activation markers and cytokines do not differ between groups, and spillover of CD20 signal into the TNFα channel differs between staining runs. These results link sensitized tolerance to an expanded dendritic-cell compartment and show that staining batches must be controlled in mass cytometry studies of food allergy.

## Introduction

IgE-mediated food allergy affects up to one in ten infants in population-based studies that confirm allergy by oral food challenge^1^, and peanut allergy usually persists^2,3^. Not every infant who produces peanut-specific IgE reacts on ingestion: a substantial proportion are sensitized but tolerant^4,5^.

Immune features that separate sensitized tolerance from clinical allergy would aid diagnosis and point to mechanisms of tolerance^6^. Neeland and colleagues profiled PBMC from 36 one-year-old infants of the population-based HealthNuts cohort^7^—12 PA, 12 PST and 12 NA infants, all classified by oral food challenge—by mass cytometry^8^ after culture in medium, peanut protein or PMA/ionomycin^6^. In unstimulated cells they reported more CD19^hi^HLA-DR^hi^ B cells in PA infants and fewer naive CD4 T cells and more pDCs in PST infants. Their unsupervised analysis applied FlowSOM^9^ to random subsamples of 50,000 cells per file, aligned marker intensities between samples by landmark registration^10^, modeled batch as a random effect, excluded clusters below 1% of cells and assessed rare populations such as pDCs by manual gating^6^.

Each of these steps is a choice. The clustering algorithm and its resolution^11,12^, transformation and subsampling^13–15^, and the handling of technical variation between samples^16–20^ can each change which populations are found and how they differ between groups^21,22^. Independent reanalysis of public data is the most direct test of whether a reported signature survives such choices^23,24^. Graph-based clustering and fast embeddings now make it practical to analyze every acquired event rather than a subsample^25,26^, but whether this improves inference in a typical clinical cohort has rarely been examined.

We reanalyzed the unstimulated condition of the published dataset on cytowhiz.ai, a platform that processes all events, to identify cell populations that differ between PA, PST and NA infants. We combined transparent cluster annotation, explicit staining-batch diagnostics and FDR-controlled comparisons of cell-type frequencies and of activation and cytokine markers, and asked whether the published differences are reproduced and, where they are not, why.

## Results

### All-event clustering resolves 40 PhenoGraph clusters, including rare populations

After quality control and gating, 3,560,317 cells were retained (median 90,946 per infant, range 21,013–207,420; Extended Data Fig. 1b). PhenoGraph partitioned them into 40 clusters ranging from 53 cells to 13.9% of all cells (Fig. 1a). Ward clustering of the median profiles separated the major lineages (cophenetic correlation 0.67), with the best mean silhouette width at k = 7 (0.35; Extended Data Fig. 1a). Rare populations formed distinct clusters without manual gating: pDCs (CD123^+^HLA-DR^+^CD4^+^CD11c^−^; 2,710 cells, 0.08%)^27^ and lineage-negative CD127^+^CD25^+^ ILC-like cells (6,488 cells, 0.18%)^28^ (Fig. 1). Clusters with similar lineage-marker profiles lay closer together on the t-SNE map (Mantel test, Spearman r = 0.33, permutation *P* = 0.0002). A cluster’s nearest neighbor on the map, however, belonged to the same major lineage only modestly more often than expected by chance (15 of 35 clusters, 43%, against 30%; *P* = 0.08), although T-cell clusters did so in 11 of 17 (65%, against 47%). The map therefore conveys the broad organization of lineages but not which cluster is most similar to which.

**Figure 1.**
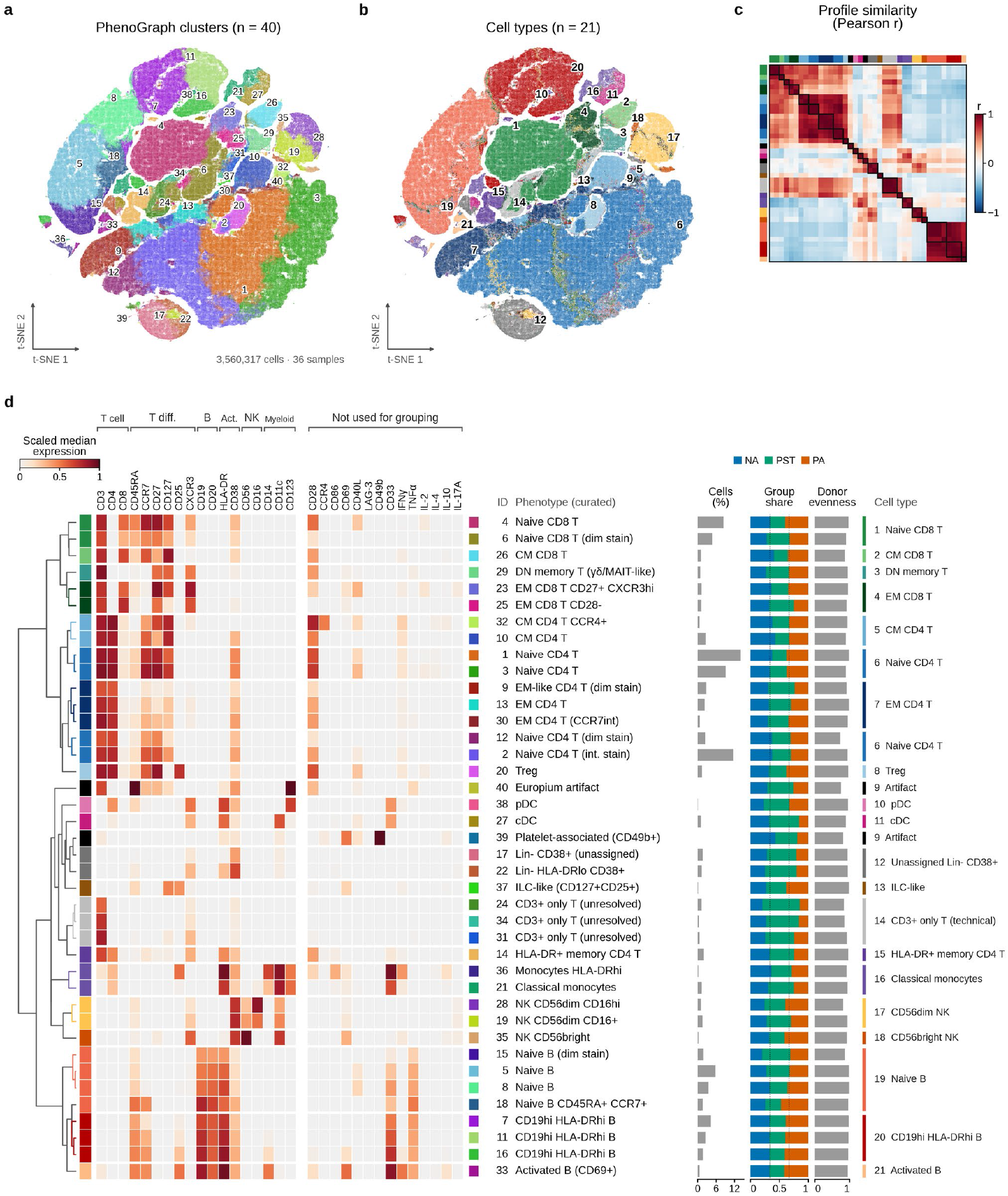
Marker-expression profiles and curated annotation of 40 PhenoGraph clusters from 3,560,317 cells. **a**, t-SNE map of all live single cells from 36 infants colored by PhenoGraph cluster; numbers mark each cluster’s densest region. All cells are shown, with opacity scaled to log density. **b**, The same map colored by curated cell type (numbers refer to the list in **d**). **c**, Pearson correlation between cluster median profiles (18 lineage markers), ordered as in **d**; boxes outline contiguous cell-type blocks. **d**, Median arcsinh-transformed (cofactor 5) expression per cluster, scaled to 0–1 between the 1st and 99.9th percentiles of all cells (minimum span 2.5 arcsinh units). Rows are ordered by Ward clustering of the 18 lineage markers; the right-hand block shows the 14 other antibody channels used by PhenoGraph but not for grouping. Row labels give the cluster number and curated phenotype; bars show cluster size, relative abundance in each clinical group (mean per-infant frequency, normalized to sum to 1; dotted lines mark equal representation) and donor evenness (normalized Shannon entropy across infants). CD33 signal in B cells reflects HLA-DR, and TNFα signal reflects CD20 (spillover within gadolinium and samarium); CD25, IFNγ, IL-10 and IL-17A carry small oxide contributions (see Results).

### Curated annotation resolves weakly stained naive T cells and technical clusters

Rule-based labeling called the three largest CD4 T-cell clusters (1, 2 and 3; 34.6% of cells) central memory, because CD45RA stained weakly in this panel (raw median arcsinh value 1.0 or below in these clusters). These cells were CCR7^+^CD27^+^CD28^+^CD38^+^ and were re-annotated as naive CD4 T cells. Together with a dim-stained copy (cluster 12) they comprised a median of 36.4% of cells per infant, against 35.8% in the published analysis. Curation yielded 19 cell types (Fig. 1b,d; Supplementary Table 1), including CCR4^+^ central memory CD4 T cells, two effector memory CD8 T-cell subsets (CD27^+^CXCR3^hi^ and CD28^−^), double-negative memory T cells with a γδ-or MAIT-like CD27^+^CD28^+^CD127^+^ phenotype, CD56^dim^ and CD56^bright^ NK cells and CD69^+^ activated B cells.

Three groups of clusters were technical: T cells that had lost every marker except CD3 (clusters 24, 31 and 34); 268 CD49b^+^CD69^+^CD40L^+^ events consistent with platelets attached to nucleated cells (cluster 39); and 53 events with equal, very bright ^151^Eu and ^153^Eu signal, the signature of natural-abundance europium contamination (cluster 40). Two channels received a fixed fraction of another channel’s signal through spillover. CD33 (^158^Gd) received 3.5–7.3% of the HLA-DR (^157^Gd) signal in every B-cell cluster (median 5%; coefficient of variation 21%, against 78% relative to CD19), spillover by one mass unit within gadolinium^29,30^. B cells therefore appeared CD33^+^ in proportion to their HLA-DR intensity (Fig. 1d), and CD33 positivity does not indicate a myeloid identity in this panel. The original investigators also excluded CD33 from their analyses because of a technical problem with its metal tag^31^. The TNFα channel (^152^Sm) received spillover from CD20 (^147^Sm) that differed between staining runs (described below). Oxide spillover (M+16) was small. IFNγ received about 1% of the CD4 signal (1–3 counts in CD4 T-cell clusters), and CD25 0–0.5% of HLA-DR in most HLA-DR^+^ clusters; only in the two clusters with the brightest HLA-DR did CD25 reach about 1% of HLA-DR (38–40 counts), which may include CD25 on activated cells or doublets. IL-10 (from CD28) and IL-17A (from CD38) received less than 1 count. CXCR3 followed known CXCR3^+^ populations (effector memory CD8 T cells, CD56^bright^ NK cells and pDCs) and showed no spillover. None of these signals was used for annotation. Two clusters (17 and 22) carried CD38 without any lineage-defining marker (cluster 22 was also HLA-DR^lo^ and CXCR3^+^) and could not be identified with this panel.

### A sample-level staining batch effect splits populations into bright and dim clusters

Twelve samples—P08–P10, P17–P22 and P32–P34—had 32.6–57.5% of their T cells in dim-stained clusters, compared with 3.6–14.7% in the other 24 samples, with no intermediate values (Fig. 2a). The same samples were dim in the B-cell lineage (ρ = 0.86, *P* = 1.3 × 10^−11^; Fig. 2b), and dimness was unrelated to the number of cells recovered (ρ = 0.05). The dim samples fell in contiguous identifier blocks, as expected for barcoding or staining batches, and came disproportionately from PST infants (6 of 12, against 3 of 12 NA and 3 of 12 PA infants).

**Figure 2.**
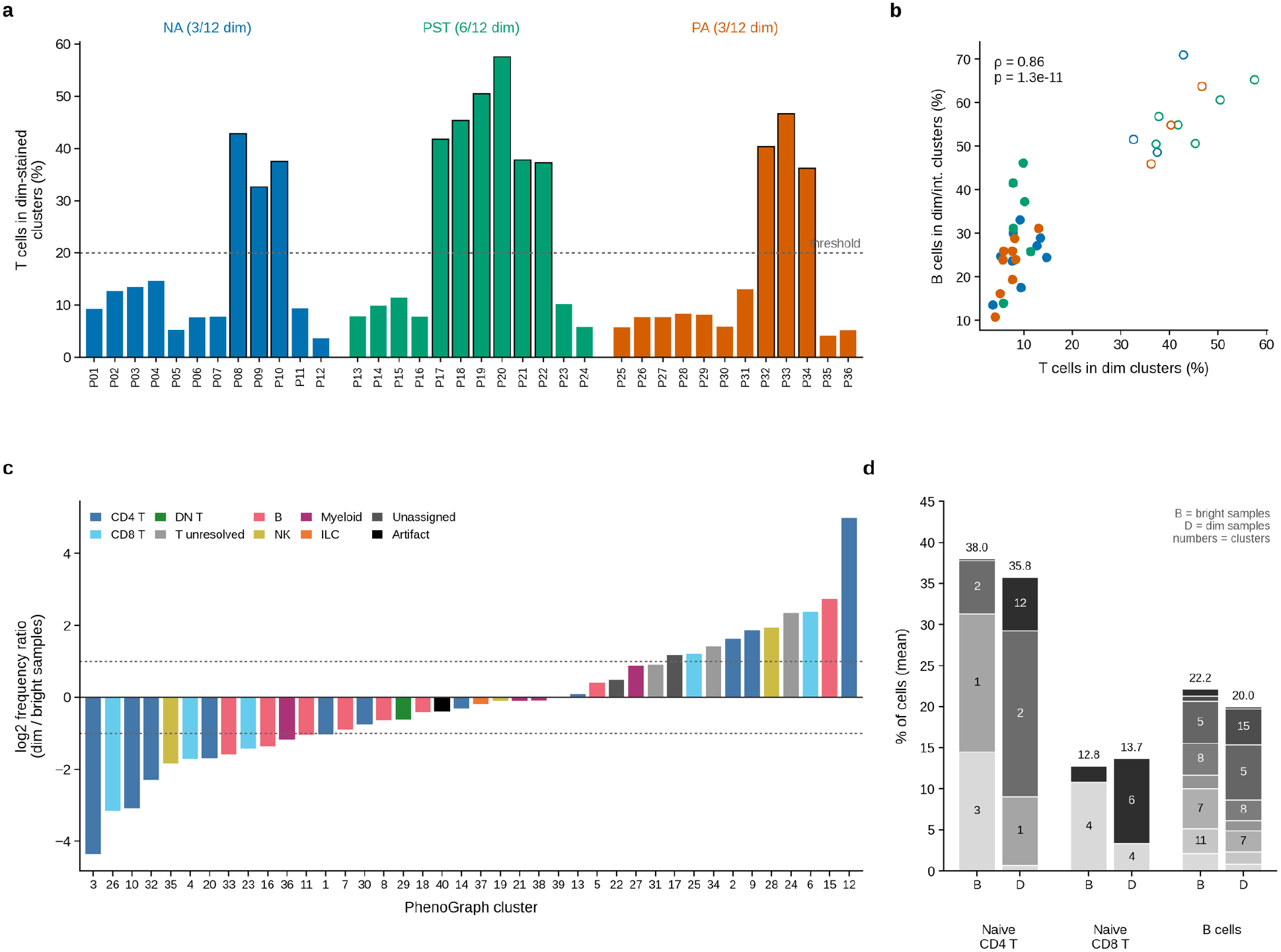
A sample-level staining batch effect. **a**, Percentage of each sample’s T cells that fall into dim-stained or CD3-only clusters; outlined bars mark dim-batch samples (above the 20% threshold, dashed line). **b**, Dim shares in T and B cells are correlated across samples (Spearman ρ); open symbols denote dim-batch samples and colors the clinical group. **c**, Ratio of each cluster’s mean frequency in dim-versus bright-batch samples (log2), colored by lineage; dashed lines mark two-fold differences. **d**, Mean composition of naive CD4 T, naive CD8 T and B cells in bright-batch (B) and dim-batch (D) samples: totals are preserved while cells move between bright and dim clusters (numbers inside bars are PhenoGraph clusters).

The effect changed how cells were assigned to clusters rather than how many cells each lineage contained. Individual clusters differed up to 32-fold in frequency between dim-and bright-batch samples (Fig. 2c), whereas lineage totals did not. Naive CD4 T cells, for example, made up 38.0% of cells in bright-and 35.8% in dim-batch samples, but cluster 3 accounted for 14.5% versus 0.7% and cluster 12 for 0.2% versus 6.5% (Fig. 2d). In the most affected sample (P20), 25.5% of T cells had lost every marker except CD3. Comparisons at the level of individual clusters were therefore confounded by batch, and all further tests used cell types that merge bright and dim variants, with batch as a covariate.

### HLA-DR^+^ memory CD4 T cells, cDCs and lineage-negative CD38^+^ cells are more frequent in PST than PA infants

No lineage total (T, CD4 T, CD8 T, B and NK cells, monocytes or dendritic cells) differed between groups (all overall *q* ≥ 0.10; Fig. 3a; Supplementary Table 2). Among the 19 cell types, the overall test was significant for lineage-negative CD38^+^ cells (*q* = 0.009) and HLA-DR^+^ memory CD4 T cells (*q* = 0.045), with a trend for conventional dendritic cells (cDC; *q* = 0.086). Of the 57 pairwise comparisons, two passed FDR control at 5% (Fig. 3b): lineage-negative CD38^+^ cells (4.5% vs 1.5% of cells; 2.26-fold, 95% CI 1.55–3.29; *q* = 0.006) and HLA-DR^+^ memory CD4 T cells (2.5% vs 1.6%; 1.97-fold, 95% CI 1.34–2.91; *q* = 0.033) were both more frequent in PST than PA infants. cDCs showed the same pattern (1.0% vs 0.4%; 2.55-fold, 95% CI 1.39–4.69; *q* = 0.069). For all three, PA infants had the lowest and PST infants the highest frequencies, with NA infants in between (PST vs NA 1.45-, 1.66-and 1.69-fold; *q* ≥ 0.14).

**Figure 3.**
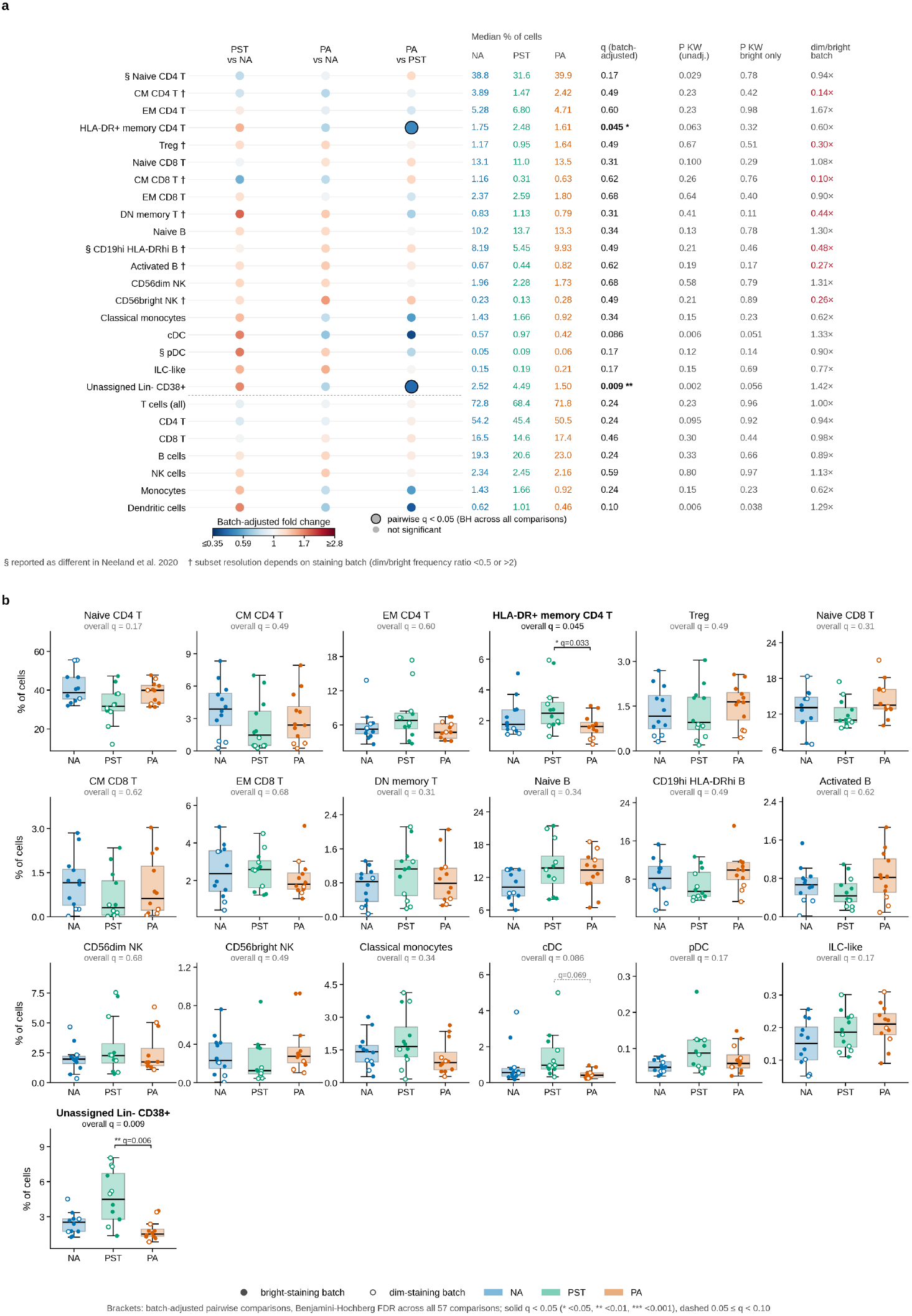
Cell-type abundance in NA, PST and PA infants. **a**, Batch-adjusted fold changes for the three pairwise comparisons (color; outlined dots, pairwise *q* < 0.05 with Benjamini–Hochberg correction across all 57 comparisons), median frequencies per group, overall *q* (F-test of group in the batch-adjusted model, Benjamini–Hochberg across 19 cell types or 7 lineage totals), Kruskal– Wallis *P* values (unadjusted, and bright-batch samples only) and batch fold change (dim/bright; red, more than two-fold). § marks populations reported as different by Neeland et al.^6^; † marks batch-sensitive subsets. **b**, Per-infant frequencies (% of all cells); boxes show the median and interquartile range, whiskers extend to 1.5 times the interquartile range; filled symbols are bright-batch and open symbols dim-batch samples. Brackets show batch-adjusted pairwise comparisons with *q* < 0.10 (solid, *q* < 0.05; dashed, 0.05 ≤ q < 0.10). Technical and artifact clusters are excluded.

Dunn’s test without batch adjustment supported the lineage-negative CD38^+^ (*q* = 0.026) and cDC (*q* = 0.045) contrasts but not the HLA-DR^+^ memory CD4 T-cell contrast (*q* = 0.27). Because the dim-staining batch included 6 of the 12 PST samples, we repeated the comparisons in bright-batch samples (9 NA, 6 PST and 9 PA). There, cDCs (0.82% vs 0.41%; 2.0-fold; Mann–Whitney *P* = 0.026) and lineage-negative CD38^+^ cells (3.1% vs 1.4%; 2.2-fold; *P* = 0.039) remained more frequent in PST than PA infants, whereas the HLA-DR^+^ memory CD4 T-cell difference was smaller (2.5% vs 1.8%; 1.4-fold; *P* = 0.15); it was largest among dim-batch samples (2.5% in PST vs 0.8% in PA). The published per-participant values showed the same pattern: HLA-DR^+^ effector memory CD4 T cells (2.7% vs 1.2%; *P* = 0.046) and dendritic cells (1.8% vs 0.6%; *P* = 0.037) were more frequent in PST than PA infants, and among bright-batch samples the dendritic-cell difference persisted (*P* = 0.028) but the T-cell difference did not (*P* = 0.48). Seven cell types—central memory CD4 T, regulatory T, central memory CD8 T, double-negative memory T, CD19^hi^HLA-DR^hi^ B, activated B and CD56^bright^ NK cells—were more than two-fold more or less frequent in one batch (Fig. 3a), so their resolution depends on staining intensity and their comparisons should be read with caution.

The lineage-negative CD38^+^ cells co-varied across samples with other poorly stained clusters, even within batch and group, and were twice as frequent in dim-as in bright-batch samples (median 4.0% vs 1.8%). Adjusting for continuous staining covariates reduced their PST vs NA difference from 1.66-fold to 1.34-fold (*P* = 0.15), whereas the PST vs PA difference persisted (1.90-fold, *P* = 0.002). The published analysis likewise assigned more cells to its undefined cluster in PST than in PA infants (7.3% vs 2.8%; *P* = 0.005). One of the two clusters was scattered across 32 separate fragments of the t-SNE map at the edge of CD4 T-cell territory, unlike genuine rare populations. These events passed DNA, event-length and viability gates, which excludes debris and non-nucleated events. They did not have a neutrophil phenotype, although neutrophils are the granulocytes that can co-purify with PBMC as low-density cells: of the neutrophil markers in the panel, CD16, CD11c and CD33 were negative in both clusters (at most 2.4 counts, against 441 counts of CD16 on CD16^hi^ NK cells and 767 counts of CD11c on classical monocytes), and one cluster carried CD38 (77 counts), which resting neutrophils express only weakly. The only CD16-bright clusters in the dataset were CD56^dim^ NK cells with the high CD38 typical of infant NK cells. Immature low-density granulocytes, which carry little CD16, cannot be excluded, because the panel lacks CD45, CD15, CD66b and progenitor markers.

### Activation markers and cytokines do not differ between groups

We next asked whether the activation state of any cell type differed between groups. For each of the 19 cell types we compared 14 functional channels (CD69, CD40L, HLA-DR, CD25, CD38, CD86, LAG-3, CD49b, IFNγ, TNFα, IL-2, IL-4, IL-10 and IL-17A), each measured per infant as the percentage of positive cells and as median intensity after spillover compensation (Methods; Extended Data Fig. 2). CD86 was stained only in samples from the first three of six acquisition batches, because the antibody stock was depleted during processing^31^, so CD86 comparisons combine stained and unstained samples. The cells had been cultured for 24 h in medium, with brefeldin A for the final 4 h^6^, so cytokine signal reflects spontaneous production in culture. That signal was sparse: IL-4 and LAG-3 were positive in fewer than 0.01% of all cells, the median percentage of IL-2^+^ and IL-17A^+^ cells stayed below 0.6% in every cell type, and 83 of 255 cell-type– marker combinations had too few positive cells to test.

No functional marker differed between NA, PST and PA infants. None of the 172 testable comparisons of the percentage of positive cells, and none of the 142 comparisons of median intensity, reached *q* < 0.10 (lowest pairwise *q* = 0.38 and 0.82), and the analysis without compensation or technical adjustment agreed (lowest *q* = 0.66). Because comparisons on the same infants are correlated, we also counted nominally significant comparisons (*P* < 0.05) and compared the count with that obtained after relabeling whole infants at random. Overall it did not exceed chance (32 against a permutation median of 23; *P* = 0.25). The one borderline excess, in the percentage of positive cells between PST and PA infants (20 against 7; *P* = 0.06), had the lower value in PST infants in 17 of the 20 comparisons across unrelated markers and cell types, the pattern expected from the dimmer staining of half of the PST samples rather than from cell-type-specific biology.

The two clusters with the brightest HLA-DR, CD69^+^ B cells and HLA-DR^hi^ monocytes, were positive for several unrelated intracellular channels at once and carried signal from other lineages (CD14, CD3 and CD28 in the B cells; CD3 and CD28 in the monocytes). They probably contain doublets or conjugates, and we did not interpret their cytokine content.

### Spillover from CD20 into the TNFα channel differs between staining runs

One signal did not fit spontaneous cytokine production. Without compensation, a fifth to a quarter of B cells read as TNFα^+^ (median 20% of naive B cells and 27% of CD19^hi^HLA-DR^hi^ B cells), whereas classical monocytes, the main source of spontaneous TNFα in culture, were 0.3% positive. A screen of every channel as a possible source for every functional channel (235 testable pairs; Methods) flagged a single pair: spillover from the CD20 channel (^147^Sm) into the TNFα channel (^152^Sm) (Extended Data Fig. 3). In cluster medians pooled across infants, the spillover coefficient was 21% in every B-cell cluster (range 18–25%; coefficient of variation 13%), whereas the ratios to HLA-DR and CD19 varied 38-and 6-fold (coefficients of variation 101% and 62%), and TNFα stayed at or below about 1 count in every cluster without CD20. Within the eight CD20^+^ clusters (787,224 cells), the two channels covaried cell by cell (Spearman ρ = 0.69; Extended Data Fig. 3a,b).

The spillover coefficient was not the same in every infant (Extended Data Fig. 3d). It was 21.8– 29.6% (median 26.2%) in 30 infants and 4.9–5.9% (median 5.5%) in the other six, with no infant in between. Within each infant it was constant across B-cell clusters (median coefficient of variation 5%), and it did not differ between clinical groups (Kruskal–Wallis *P* = 0.90) or, among the 30, between dim-and bright-batch samples (Mann–Whitney *P* = 0.66). The six (P11, P12, P23, P24, P35 and P36) were the last two identifiers of each group, consistent with a separate staining run. A pooled coefficient therefore could not compensate every sample: subtracting the pooled 21% reduced TNFα^+^ cells in CD20^+^ clusters from 26.0% to 5.4%, but it over-corrected the six infants and left 1.8–10.8% TNFα^+^ naive B cells in the other 30. Per-infant coefficients reduced TNFα^+^ cells in CD20^+^ clusters to 2.7% (Extended Data Fig. 3c,e). The remainder was highest in the infants with the strongest uncompensated signal, as expected from counting noise when a large spillover is subtracted cell by cell, although a small genuine TNFα^+^ fraction cannot be excluded. The spillover thus raises the detection limit for TNFα in B cells.

Signal at a heavier isotope of the same element can arise from isotopic impurity of an enriched metal, but rarely at more than a few percent^29^, far below the 26% seen in most infants. Natural-abundance samarium is also excluded, because it would have produced signal in the other channels at samarium masses (144, 149, 150 and 154), which B cells lacked. The likeliest source is therefore the reagent, for example ^152^Sm carried by the CD20 conjugate, with a different conjugate or cocktail preparation in one staining run. Uncompensated, the spillover made 4.7% of all analyzed cells TNFα^+^ at our threshold, 99% of them B cells (fewer than 0.1% of non-B cells were positive), so that the frequency of TNFα^+^ cells tracked each infant’s B-cell frequency (Spearman ρ = 0.44, *P* = 0.007).

### The published naive CD4 T-cell and B-cell differences track the staining batch

The digitized per-participant values reproduced the published statistics. CD19^hi^HLA-DR^hi^ B cells (published B-cell cluster 2) were more frequent in PA than PST infants (Mann–Whitney *P* = 0.005; published *P* = 0.001) and NA infants (*P* = 0.038; published *P* = 0.01), and naive CD4 T cells were less frequent in PST than NA infants (*P* = 0.010; published *P* = 0.006). Median frequencies of 14 matched populations agreed closely between the analyses (ρ = 0.86; Fig. 4a)—for example, naive CD4 T cells 36.4% vs 35.8%, B cells 21.9% vs 19.4% and HLA-DR^+^ memory CD4 T cells 1.8% vs 1.7%. Per-infant agreement was high for most populations (B cells ρ = 0.99, naive CD8 T cells 0.93, naive CD4 T cells 0.89, HLA-DR^+^ memory CD4 T cells 0.86) and lower for small ones (dendritic cells 0.68, monocytes 0.35). Composition fingerprints confirmed the sample mapping: for 21 of the 36 infants the published participant with the same identifier was the closest of 36 candidates, and for 32 it was among the three closest (median rank 1, against 18 by chance; none of 2,000 permutations was as good, *P* < 0.0005).

**Figure 4.**
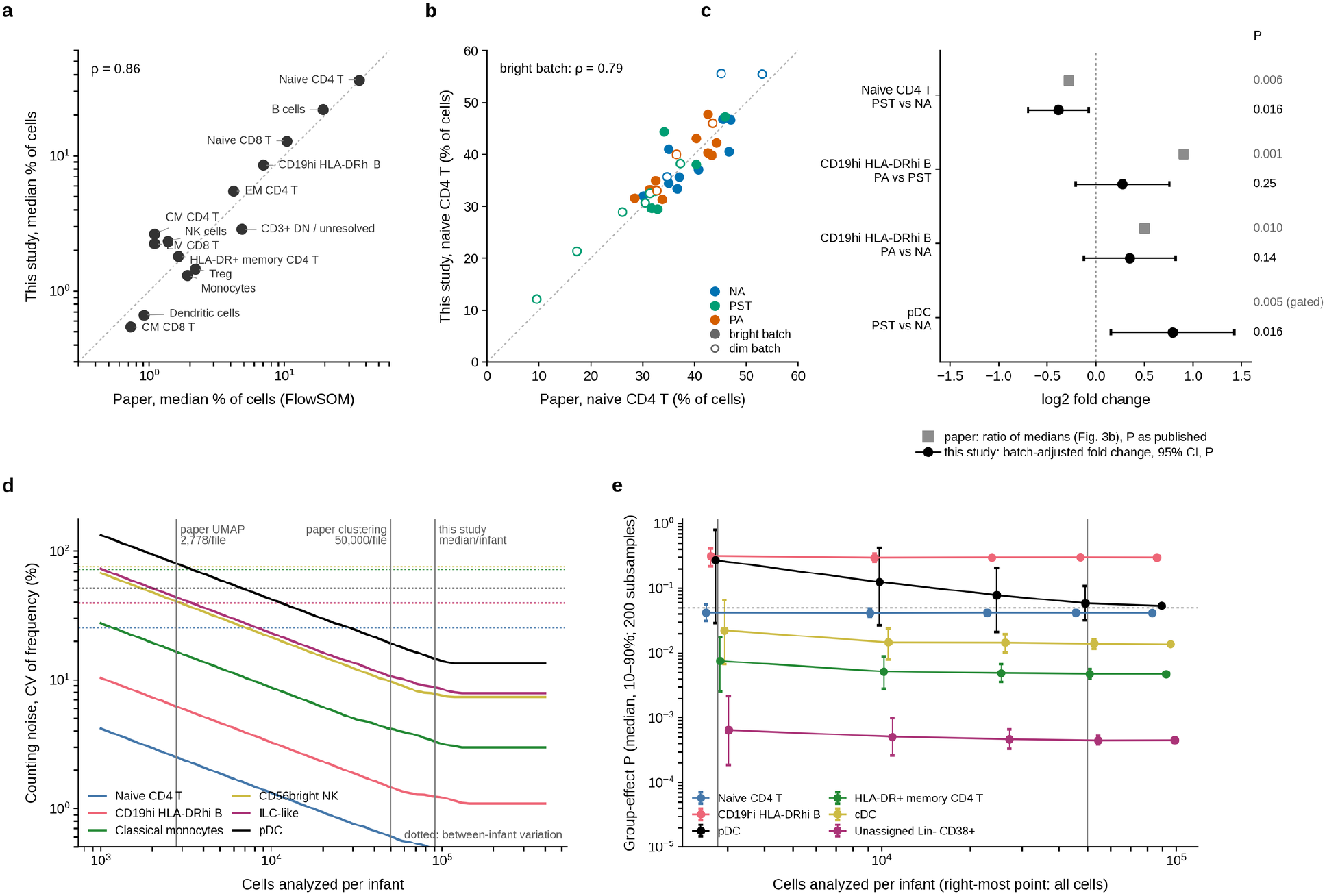
Concordance with the published analysis and the effect of the number of analyzed events. **a**, Median frequencies of 14 matched populations (log scale) in the published FlowSOM analysis^6^ (values digitized from its Fig. 3b) and in this study. **b**, _Per-infant naive CD4 T-cell frequencies; colors denote clinical group and open symbols dim-batch samples; ρ, Spearman correlation in bright-batch samples. **c**, The published signatures: ratio of group medians in the published per-participant values (gray squares; published *P* values, pDC from manual gating) and batch-adjusted fold change with 95% confidence interval in this study (black; *P* values). **d**, Counting noise (coefficient of variation of a frequency estimate) as a function of cells analyzed per infant for six populations, against between-infant variation (dotted lines); vertical lines mark 2,778 and 50,000 cells per file (published UMAP and clustering subsamples) and the median number of cells per infant in this study. **e**, Group-effect *P* values (median and 10–90% range of 200 subsamples) after subsampling each infant to the indicated number of cells; right-most points use all cells._

Of the three published signatures (Table 1; Fig. 4c), only the pDC increase was independent of staining batch. pDCs were more frequent in PST than NA infants (batch-adjusted fold change 1.73, 95% CI 1.11–2.69; *P* = 0.016; *q* = 0.15), including among bright-batch samples (2.0-fold; Mann– Whitney *P* = 0.036). Naive CD4 T cells were less frequent in PST than NA infants across all samples (0.76, 0.62–0.95; *P* = 0.016; *q* = 0.15) but not among bright-batch samples (0.96-fold; *P* = 0.61). CD19^hi^HLA-DR^hi^ B cells were not significantly more frequent in PA than PST infants (1.21-fold, 0.87– 1.69; *P* = 0.25; bright-batch samples *P* = 0.27) or NA infants (1.27-fold, 0.92–1.76; *P* = 0.14). The published per-participant values behaved the same way: among bright-batch samples, the naive CD4 T-cell difference between PST and NA infants fell from *P* = 0.010 to *P* = 0.22, and the B-cell difference between PA and PST infants from *P* = 0.005 to *P* = 0.13 (Table 1).

**Table 1.** Re-evaluation of the published abundance signatures in unstimulated PBMC. Published *P* values are from χ^2^ tests in mixed-effects models (FlowSOM clusters) or manual gating^6^. Recomputed values are Mann–Whitney tests on per-participant frequencies digitized from the published Fig. 3b. Fold changes compare the first with the second group named. Values after the slash and in parentheses are Mann–Whitney tests restricted to bright-batch samples (9 NA, 6 PST and 9 PA).

| Published signature | Published result<sup>6</sup> | Recomputed from published values: all / bright-batch samples | This study: group medians (% of cells) | Batch-adjusted fold change (95% CI) | $P$ (bright-batch samples) | $q$ (57 comparisons) |
| --- | --- | --- | --- | --- | --- | --- |
| Naive CD4 T cells lower in PST than NA | $P = 0.006$ (FlowSOM; omnibus 0.007) | $P = 0.010 / 0.22$ | 31.57 vs 38.80 | 0.76 (0.62–0.95) | 0.016 (0.61) | 0.15 |
| CD19<sup>hi</sup>HLA-DR<sup>hi</sup> B cells higher in PA than PST | $P = 0.001$ (FlowSOM; omnibus 0.005) | $P = 0.005 / 0.13$ | 9.93 vs 5.45 | 1.21 (0.87–1.69) | 0.25 (0.27) | 0.50 |
| CD19<sup>hi</sup>HLA-DR<sup>hi</sup> B cells higher in PA than NA | $P = 0.01$ (FlowSOM) | $P = 0.038 / 0.10$ | 9.93 vs 8.19 | 1.27 (0.92–1.76) | 0.14 (0.60) | 0.40 |
| pDCs higher in PST than NA | $P = 0.005$ (manual gating) | not in Fig. 3b | 0.088 vs 0.048 | 1.73 (1.11–2.69) | 0.016 (0.036) | 0.15 |

Both populations depend on staining intensity, and the dim batch included half of the PST samples. In dim-batch samples, a third to more than half of T cells fell into dim-stained clusters, and in some samples many had lost CD4 staining. The published analysis placed CD3^+^ cells lacking CD4 and CD8 in two undefined clusters and excluded them from testing^6^; in the published values these clusters held more cells in dim-batch than in bright-batch samples (median 6.7% vs 4.0%; *P* = 0.002), and their size tracked the share of dim-stained T cells measured here (ρ = 0.80, *P* = 3.7 × 10^−9^). The lowest published naive CD4 T-cell frequencies were in two dim-batch PST samples, P20 (9.5%) and P22 (17.2%), where our pipeline also measured low values (12.1% and 21.3%; Fig. 4b), and 22.9% of the cells of P20 were in the published undefined T-cell clusters. Likewise, the published CD19^++^HLA-DR^++^ B-cell cluster made up 4.8% of cells in dim-batch samples against 8.2% in bright-batch samples (*P* = 0.003), and our CD19^hi^HLA-DR^hi^ B cells were half as frequent in dim-batch samples. Because the dim samples came disproportionately from PST infants, both populations appear reduced in PST infants. The published analysis also assigned 10–11% of cells to dendritic cells in two dim-batch samples, P08 and P22; we found 4.0% and 5.1%, against a median of 0.7%.

### Event number does not explain the differences from the published analysis

None of the differences between the analyses depends on the number of events analyzed. Naive CD4 T cells and CD19^hi^HLA-DR^hi^ B cells are abundant, so at the 50,000 cells per sample used for the published clustering their frequencies carry counting errors of about 0.6% and 1.5%, against between-infant variation of 25% and 39% (Fig. 4d); for pDCs, the rarest population tested, the counting error was 19% against 52%. The published pDC result came from manual gating rather than the subsample^6^. Conversely, the published per-participant values already contained the PST– PA differences in dendritic cells (*P* = 0.037) and HLA-DR^+^ effector memory CD4 T cells (*P* = 0.046) reported here; they were not tested pairwise because their three-group tests did not pass FDR control^6^. Within our data, subsampling each infant to 50,000 cells reproduced every full-data *P* value (Fig. 4e; pDC median *P* 0.059 vs 0.053; HLA-DR^+^ memory CD4 T cells 0.0048 vs 0.0047). Event number mattered only at 2,778 cells per infant—the subsample of the published UMAP—where counting noise for pDCs rose to 80% and their difference was lost (median *P* 0.27). Analyzing all events also revealed technical events present at a few dozen to a few hundred cells in the whole dataset (53 europium-contaminated and 268 platelet-associated events) and allowed staining failure to be quantified in individual samples. Because subsampling reused the full-analysis clusters, it does not test whether clustering fewer events would have resolved the same rare populations.

## Discussion

Three cell populations were more frequent in peanut-sensitized tolerant than in peanut-allergic infants: HLA-DR^+^ memory CD4 T cells, conventional dendritic cells and lineage-negative CD38^+^ cells, with non-allergic infants in between. Of the three published signatures, the increase of plasmacytoid dendritic cells in tolerant infants was reproduced, whereas the naive CD4 T-cell and B-cell differences tracked a staining batch. Activation markers and cytokines did not differ between groups, and population frequencies agreed closely with the original analysis, infant by infant.

The dendritic-cell findings were the most robust. cDCs were more frequent in tolerant than allergic infants in both analyses and among bright-batch samples, and pDCs were more frequent in tolerant than non-allergic infants, as originally reported^6^. Plasmacytoid dendritic cells contribute to tolerance to innocuous antigens^27^, so an expanded dendritic-cell compartment in sensitized-tolerant infants is plausible, although this panel cannot assess dendritic-cell function. HLA-DR^+^ memory CD4 T cells, a population whose frequency matched the published analysis closely, were twice as frequent in PST as in PA infants in both analyses, but the difference was concentrated in dim-batch samples and small among bright-batch ones, so it is the least robust of the three. HLA-DR appears on T cells days after activation, so the difference probably reflects a state that predates the food challenge 2 h before sampling, although activation signals received during the challenge could have been expressed during culture. The lineage-negative CD38^+^ cells were also more frequent in PST infants, but their abundance tracked staining quality even within batch, the published analysis assigned more cells to its undefined cluster in PST infants, and adjustment for continuous staining covariates removed their difference from NA infants, though not from PA infants. They lacked the neutrophil markers the panel contains (CD16, CD11c and CD33) and expressed CD38, so we regard them as largely poorly stained nucleated cells. Immature low-density granulocytes, which carry little CD16, remain the main biological alternative; CD45, CD15, CD66b and CD34 would resolve their identity.

The main technical finding was a sample-level staining batch that affected a third of the samples, half of them from sensitized-tolerant infants. Three features identify it as technical: the dim samples form contiguous identifier blocks, the dimming is coherent across T-and B-cell markers, and it changes how cells are assigned to clusters without changing lineage totals. In un-normalized data, any clustering algorithm will split a population by intensity when intensity differs systematically between samples, producing bright and dim duplicates whose frequencies track batch rather than biology^17,18^. The original authors anticipated this and registered marker intensities across samples before clustering^6,10^, but CD45RA and CD38 were not registered for the unstimulated samples^6^. Weak CD45RA staining in dim-batch samples would push naive T cells toward memory or unclassified clusters, consistent with the low naive CD4 T-cell frequencies that both analyses found in P20 and P22 and with the high dendritic-cell frequencies in P08 and P22. Whether the batch reflects staining or acquisition sensitivity can be decided from the DNA-intercalator signal, which does not depend on antibody staining. If it is also dimmer in these samples, global intensity scaling or bead-based normalization would correct it^32^; a marker-specific deficit requires per-channel normalization such as landmark registration, CytoNorm, cyCombine or CytofRUV^10,18–20^.

Of the three published signatures, only the pDC increase was independent of staining batch. The naive CD4 T-cell and B-cell differences concern populations defined by staining intensity, and neither held among bright-batch samples, in the published values or in ours. In the published analysis, T cells that had lost CD4 staining fell into undefined clusters that were excluded from testing, which lowers naive CD4 T-cell frequencies in exactly the samples where staining failed. The manual gating used to confirm these results^6^ depends on the same CD4, CD45RA, CD19 and HLA-DR intensities. Because half of the PST samples were in the dim-staining batch, findings specific to this group are the most exposed to staining artifacts. Reporting per-sample staining diagnostics, batch membership and analyses that exclude batch-affected samples would make such findings easier to evaluate^12,16^.

Activation markers and cytokines did not differ between groups in unstimulated culture, even though blood was drawn 2 h after a challenge that only PA infants failed. Functional differences therefore have to be sought in the responses to stimulation, where the original study reported them^6^, although 24 h of culture may also have masked activation present in vivo. Two technical findings bear on that analysis.

First, spillover put about a quarter of the CD20 signal into the TNFα channel in most infants, so B cells contribute to any count of TNFα^+^ cells that is not compensated for it. The stimulated samples were stained with the same panel, and the published increase in TNFα^+^ cells among all live cells of PA infants after PMA/ionomycin^6^ is such a count. Because the spillover coefficient differed between staining runs, it has to be estimated per sample, for example from each infant’s unstimulated sample, before TNFα is gated; doing so would show whether the spillover contributes to the published difference.

Second, CD40L, one of the two markers used to identify peanut-specific CD4 T cells^6,33^, receives spillover from CCR7 and CD27, which are bright on naive T cells. Compensating CD40L before activation-induced-marker gating would remove a background that is largest in naive cells.

The CD20 spillover was far larger than isotopic impurity produces, differed between staining runs and was found only by screening every channel pair; a spillover matrix measured once, for example with single-stained beads^29^, would not have corrected it in every sample. Such a screen is cheap and should precede any functional analysis of this panel.

The number of analyzed events accounted for none of the differences between the analyses. Counting noise falls with the square root of the number of cells and of the population frequency, and at 50,000 cells per infant it was already small relative to biological variation between infants for every population above about 0.1% of cells. The unit of replication in such studies is the participant, not the cell^21,34^, and statistical power here was limited by having 12 infants per group. The benefits of all-event processing were qualitative: rare populations such as pDCs and ILC-like cells emerged as clusters without manual gating, and technical events present at a few dozen cells per dataset became visible. Part of this benefit reflects clustering resolution—40 PhenoGraph clusters without a size cut-off, against 25 FlowSOM metaclusters with a 1% cut-off—rather than the number of events alone. Subsampling as severe as that used for the published UMAP (2,778 cells per file) would have lost the pDC signal.

This study has limitations. Batch membership was inferred from the data, not from acquisition records; the inference is supported by its bimodality and its coherence across lineages, but the original batch assignments would allow a direct test. Only six PST samples were in the bright batch, so analyses restricted to bright-batch samples cannot exclude smaller differences. The clinical groups also differed in parental country of birth and egg allergy^6^, which could not be adjusted for with 12 infants per group. Published per-participant values were digitized from a figure, with a precision of about 0.2 percentage points. Annotation involved expert judgment, documented cluster by cluster in Supplementary Table 1. Finally, only the unstimulated condition was analyzed. The published findings after peanut and PMA/ionomycin stimulation were not re-evaluated, and re-evaluating the TNFα findings will first require compensating the CD20 spillover in each sample. The unstimulated cells had been cultured for 24 h, which may have masked differences present in vivo.

In unstimulated PBMC of one-year-old infants, sensitized tolerance differed from peanut allergy in the frequencies of three cell populations—HLA-DR^+^ memory CD4 T cells, conventional dendritic cells and lineage-negative CD38^+^ cells, all higher in tolerant infants. The published increase of plasmacytoid dendritic cells in tolerant infants was reproduced, the published naive CD4 T-cell and B-cell differences tracked a staining batch, and resting activation and cytokine markers did not differ. The dendritic-cell differences were the most robust to staining quality, and the lineage-negative cells remain unidentified. Staining batch and run-dependent spillover need to be controlled explicitly, and the differences require confirmation in an independent cohort before they can be used to distinguish tolerance from allergy.

## Methods

### Study population and data

We used the mass cytometry data of the unstimulated (medium) condition from Neeland et al.^6^. The original study included 36 one-year-old HealthNuts infants^1,7^: 12 PA (objective reaction at oral food challenge with peanut skin-prick test ≥2 mm or specific IgE ≥0.35 kUA/L), 12 PST (sensitized with a negative challenge) and 12 NA (non-sensitized with a negative challenge). Blood was collected 2 h after the challenge; PBMC were isolated, cryopreserved, thawed, rested overnight and cultured for 24 h in medium with brefeldin A for the final 4 h, then palladium-barcoded^35^, stained with 24 surface and 8 intracellular metal-conjugated antibodies and acquired on a Helios mass cytometer; bead normalization^32^ and debarcoding were performed by the original investigators^6^. Per-sample FCS files were obtained from ImmPort (study accession SDY2015)^31^. Participant identifiers P01–P36 and clinical groups (P01–P12 NA, P13–P24 PST, P25–P36 PA) followed the original publication. Sample codes in the platform output were assigned to participant identifiers by upload order (NA, PA, PST; Supplementary Data) and checked against the published per-participant frequencies (see Comparison with the published analysis).

### Pre-processing and gating

Files were processed on cytowhiz.ai. Signal quality was controlled per file with PeacoQC^36^ using default settings. Events were then gated (Extended Data Fig. 4) to exclude normalization beads (^140^Ce-positive events), to retain nucleated single cells on the two DNA-intercalator channels (^191^Ir vs ^193^Ir) and on event length, and to remove dead cells (^115^In DOTA-maleimide-positive). The same gates were applied to all files, retaining 3,560,317 cells.

### Dimensionality reduction and clustering

t-SNE^37,38^ (Euclidean distance, perplexity 30, 1,000 iterations, learning rate 200) and PhenoGraph^25^ (k = 30 nearest neighbors) were computed on all 3,560,317 cells using all 32 antibody channels; the live/dead channel was not used because cells had already been gated as live. PhenoGraph returned 40 clusters.

### Cluster profiling and data-driven grouping

For profiling, intensities were arcsinh-transformed (cofactor 5) and the median of each marker was computed per cluster over all cells. Each marker was scaled to 0–1 between its 1st and 99.9th percentile across all cells, with a minimum span of 2.5 arcsinh units so that channels without a positive population (for example, cytokines in unstimulated cells) were not stretched. Clusters were grouped by Ward hierarchical clustering^39^ of the scaled profiles of the 18 lineage markers used for unsupervised clustering in the original study (CD3, CD4, CD8, CCR7, CD45RA, CD25, CD127, CD27, CXCR3, CD19, CD20, CD38, CD56, CD14, CD16, HLA-DR, CD11c and CD123)^6^, using Euclidean distances and optimal leaf ordering^40^; the number of groups was chosen by the mean silhouette width^41^ over k = 6–20. t-SNE maps show all cells.

To test whether the t-SNE map placed similar clusters together, each cluster was located at the density mode of its cells on the map. Distances between clusters on the map were compared with Euclidean distances between their scaled median profiles of the 18 lineage markers by a Mantel test (Spearman correlation; 5,000 permutations of cluster labels), and the fraction of clusters whose nearest neighbor on the map belonged to the same major lineage (T, B, NK, myeloid or other) was compared with 5,000 permutations of lineage labels. Technical and artifact clusters were excluded, leaving 35 clusters.

### Cluster annotation

Provisional phenotypes were assigned by comparing scaled cluster medians with Otsu positivity thresholds (estimated on 300,000 randomly sampled cells and constrained to 0.15–0.60 of the scaled range), following the gating hierarchy of the original study^6^. They were then curated using raw medians, marker maps on the t-SNE embedding (Extended Data Fig. 5) and the dependence of each cluster’s frequency on staining batch (next section). Because CD45RA stained weakly in this panel, naive and memory T cells were distinguished mainly by CCR7, CD27, CD28, CD127 and CD38, the last being brighter on naive than on memory T cells in these samples (Extended Data Fig. 2b). Signals that tracked the intensity of a bright channel at a neighboring mass or 16 mass units lighter—for example CD33 (^158^Gd) tracking HLA-DR (^157^Gd) in B cells—were treated as spillover rather than expression^29,30^. Clusters were assigned to 19 cell types and two technical categories (Supplementary Table 1).

### Staining-batch diagnostics

Clusters whose phenotype was a uniformly dimmer copy of a brighter cluster, or that had lost every marker except CD3, were flagged as dim-stained. For each sample we computed the percentage of T cells in these clusters, and the percentage of B cells in dim or intermediate B-cell clusters. The T-cell measure was bimodal (Fig. 2a), and samples above 20% were assigned to a dim-staining batch. For each cluster, the ratio of its mean frequency in dim-versus bright-batch samples measured its dependence on batch.

### Statistical analysis

The frequency of each cell type was expressed as a percentage of all cells per infant. The primary test was a linear model of log frequency (0.5 added to counts) on clinical group and staining batch. An F-test of the group term (2 df) gave an overall *P* value per cell type, and model contrasts gave batch-adjusted fold changes with 95% confidence intervals for PST vs NA, PA vs NA and PA vs PST. P values were adjusted with the Benjamini–Hochberg procedure^42^ across the 19 cell types for the overall test and across all 57 pairwise comparisons for the contrasts; lineage totals formed separate families of 7 and 21 tests. As sensitivity analyses we used the Kruskal–Wallis test^43^ with Dunn’s post hoc test^44^ (Benjamini–Hochberg across the 57 comparisons) without batch adjustment, and the Kruskal–Wallis test restricted to bright-batch samples (9 NA, 6 PST and 9 PA), with Mann– Whitney tests for pairwise contrasts. Technical and artifact clusters were excluded. Tests were two-sided; *q* < 0.05 was considered significant and 0.05 ≤ *q* < 0.10 a trend. For the unidentified lineage-negative CD38^+^ cells, models were additionally adjusted for continuous staining covariates (logit-transformed dim shares of T and B cells).

### Activation markers and cytokines

For every infant and cell type with at least 50 cells, we computed for each of 14 functional channels the percentage of positive cells and the median arcsinh-transformed intensity (cofactor 5). The positivity threshold of each channel was the larger of an Otsu threshold on 300,000 randomly sampled cells and 20 counts; every channel except HLA-DR (35 counts) used the 20-count floor. Markers that define a cell type (for example CD25 in Treg) were not tested. A percentage was tested if the cell type had at least 20 positive cells in total and positive cells in at least 25% of infants, and an intensity if it varied between infants.

In the primary analysis, compensated values were modeled by linear regression on group plus technical covariates: the median intensity, in the same cells, of each channel that can contribute spillover to the marker and of a lineage reference marker of the cell type (CD3, CD20, CD16, CD56, CD11c, CD123, CD127 or CD38). Percentages were arcsine-square-root transformed, and pairwise contrasts were tested with t statistics. A secondary analysis used uncompensated values with Kruskal–Wallis and Dunn’s tests^43,44^. P values were adjusted with the Benjamini–Hochberg procedure^42^ across all pairwise comparisons of a metric, and a difference was reported only if it also reached 1.5-fold and 0.5 percentage points (percentages) or 0.1 arcsinh units (intensities).

### Spillover screen and compensation

Functional channels were compensated per cell on raw counts in two steps. First, every other channel was screened as a possible source of spillover, using cluster medians pooled across infants. A pair was flagged if, across at least three clusters with source signal of at least 20 counts, the target-to-source ratio was at least 2% and constant (coefficient of variation ≤35%), the target reached at least 3 counts, and the target stayed at or below 2 counts in every cluster without source signal (below 2 counts). Spillover adds a fixed fraction of the source signal to the target channel, whereas co-expression gives ratios that differ between cell subsets. Because this fraction, the spillover coefficient, can differ between staining runs, the coefficient of a flagged pair was estimated per infant, as the median over the infant’s source-positive clusters (at least 100 cells each) of the ratio of target to source medians, and subtracted from that infant’s cells.

Second, spillover from channels one mass unit lighter or heavier (abundance sensitivity) or 16 units lighter (oxide formation)^29,30^ was subtracted, with coefficients estimated as the slope through the origin of the 25th percentile of the target across bins of source intensity (upper half of the source distribution; capped at 5%). Because the target channels are mostly background in unstimulated cells, this lower envelope is dominated by spillover. Coefficients were estimated one source at a time on the residual, largest first, so that co-expressed sources such as CCR7 and CD27 were not subtracted twice; the largest were 1.2% (CD14 into IFNγ and CCR7 into CD40L). HLA-DR and CD38 were not compensated, because their own signal far exceeds any plausible spillover. Spillover into phenotype channels (CD33) was assessed from the same cluster medians within lineages and was not compensated, because those channels were not used for annotation or testing.

### Permutation test of the number of nominal differences

Comparisons on the same infants are correlated, so the number of nominally significant comparisons expected under the null hypothesis is not simply 5% of tests. We relabeled whole infants at random 5,000 times, recomputed Dunn’s tests for all tested combinations and compared the observed number of comparisons with *P* < 0.05 with its permutation distribution.

### Comparison with the published analysis

Per-participant frequencies of the 17 FlowSOM populations of the published analysis were digitized from the stacked bars of its Fig. 3b^6^. The page was rendered at 300 dpi; each bar spans 545 pixels, so one pixel corresponds to 0.18 percentage points, and populations were assigned from the figure’s color key. Digitization was validated by re-testing the published comparisons. To verify that sample codes in the platform output mapped to the published participant identifiers, we compared the composition of five well-resolved populations (B cells, naive CD8 T cells, NK cells, monocytes and naive CD4 T cells; z-scored) between each infant and all 36 published participants, recorded the rank of the participant with the same identifier, and compared the median rank with 2,000 random permutations of identifiers.

### Number of analyzed events

Counting noise was computed as the coefficient of variation of a binomial frequency estimate, √((1 − p)/(np)), for n cells per infant (capped at each infant’s total) and population frequency p, and compared with between-infant variation, estimated as the residual standard deviation of the log-frequency model. To test whether analyzing fewer cells would have changed the conclusions, each infant’s cluster counts were subsampled without replacement (multivariate hypergeometric draws, 200 per size) to 2,778 cells (the per-file subsample of the published UMAP), 10,000, 25,000 or 50,000 cells (the published clustering subsample), keeping the cluster assignments of the full analysis, and the primary test was re-run.

### Software

Analyses used Python 3.12 with NumPy^45^, SciPy^46^, pandas^47^ and Matplotlib^48^.

## Data availability

The original FCS files are available from ImmPort (study accession SDY2015; https://doi.org/10.21430/M3VPY49EGU)^31^. Digitized per-participant values from the published Fig. 3b, cluster profiles, per-infant frequencies and functional-marker values, the spillover screen with per-infant spillover coefficients, and all test statistics are provided as Supplementary Data.

## Code availability

The analysis code (phenograph_cluster_profiles.py, celltype_abundance_vs_paper.py and functional_markers_by_group.py) will be deposited in a public repository before publication.

## Acknowledgments

We thank Neeland et al. and the HealthNuts investigators and participants for generating and sharing the data, and Symphonize for help in developing the cytowhiz.ai platform.

## Author contributions

C.C. led the analysis and helped develop the platform. K.I.A. conceptualized the study, developed the platform and directed the analysis. C.C. and K.I.A. wrote the manuscript.

---

**Extended Data Fig. 1.**
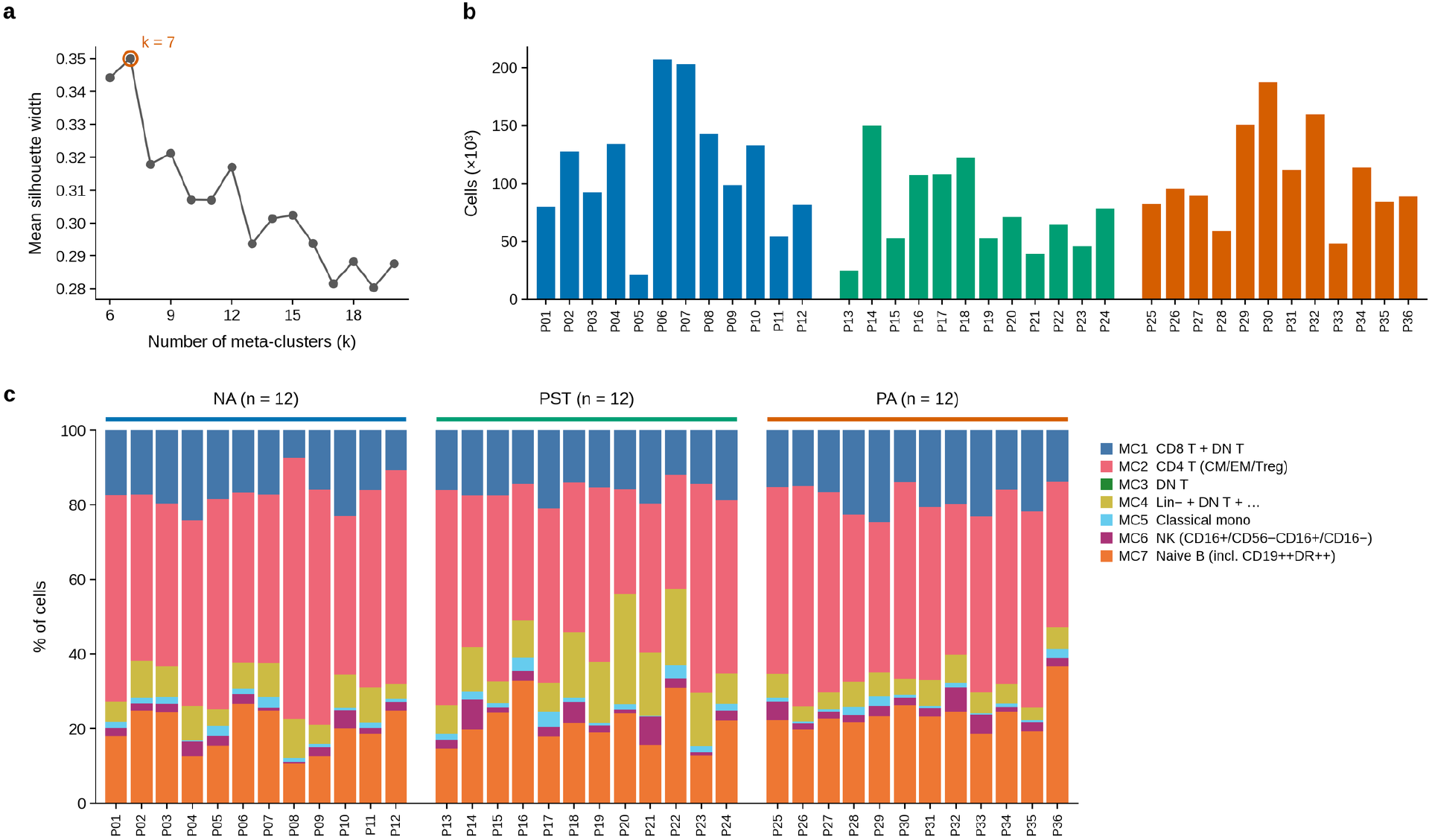
Data-driven grouping and per-infant composition. **a**, Mean silhouette width of the Ward tree cut at k = 6– 20; the maximum at k = 7 defined the data-driven meta-clusters. **b**, Cells retained per infant after quality control and gating. **c**, Per-infant composition by the seven data-driven meta-clusters (before curation).

**Extended Data Fig. 2.**
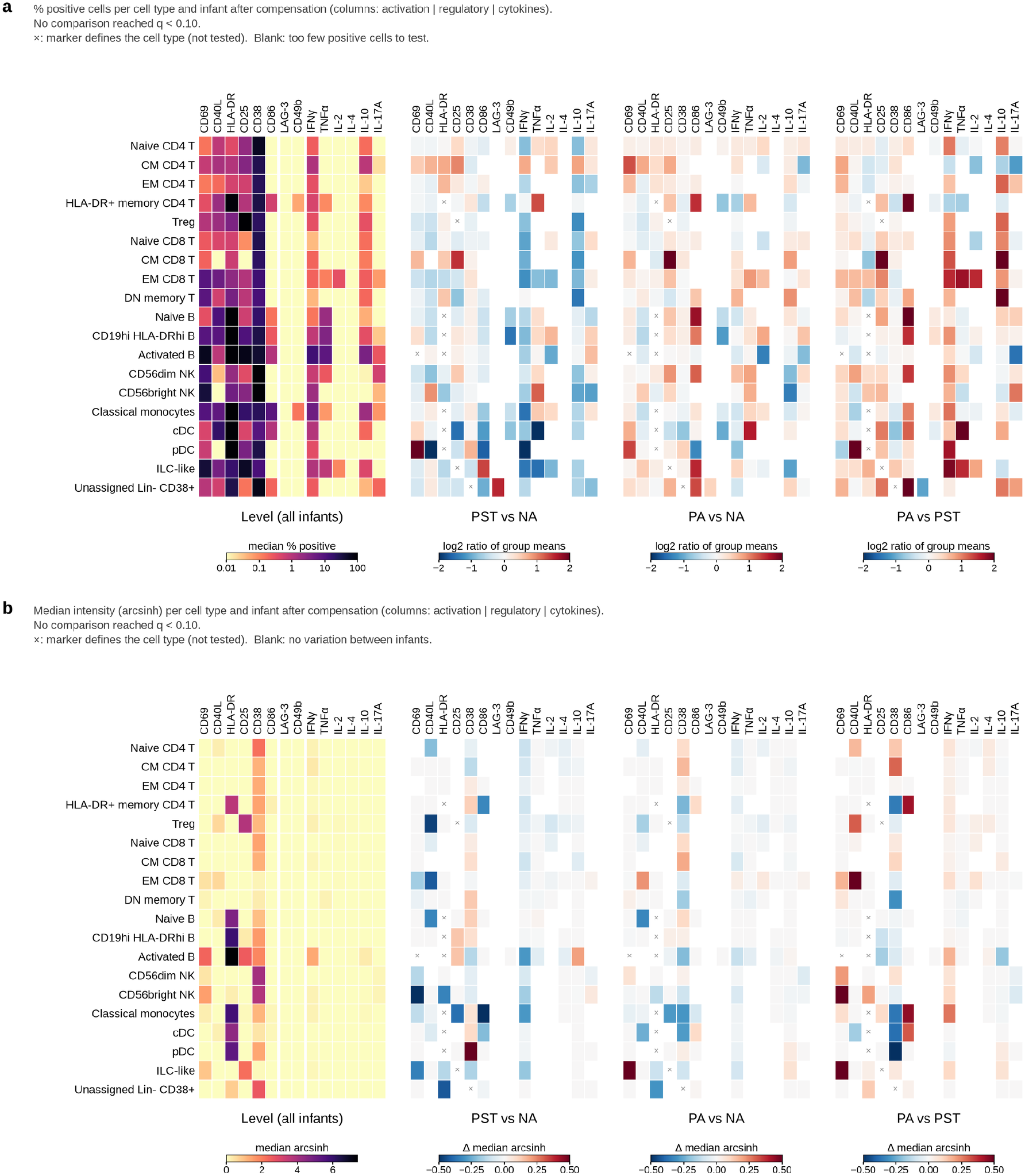
Activation markers and cytokines in unstimulated cells. **a**, Percentage of positive cells and **b**, median intensity (arcsinh) of 14 functional channels in 19 cell types after compensation; columns are grouped into activation, regulatory and cytokine markers. Left, median across all 36 infants. Right, pairwise comparisons colored by the log2 ratio of group means (**a**) or by the difference of group medians (**b**). Filled and open circles would mark comparisons with *q* < 0.05 and *q* < 0.10 in the compensated, technically adjusted model that also met the effect-size criteria (Methods); none did. ×, marker defines the cell type (not tested); blank, too few positive cells (**a**) or no variation between infants (**b**).

**Extended Data Fig. 3.**
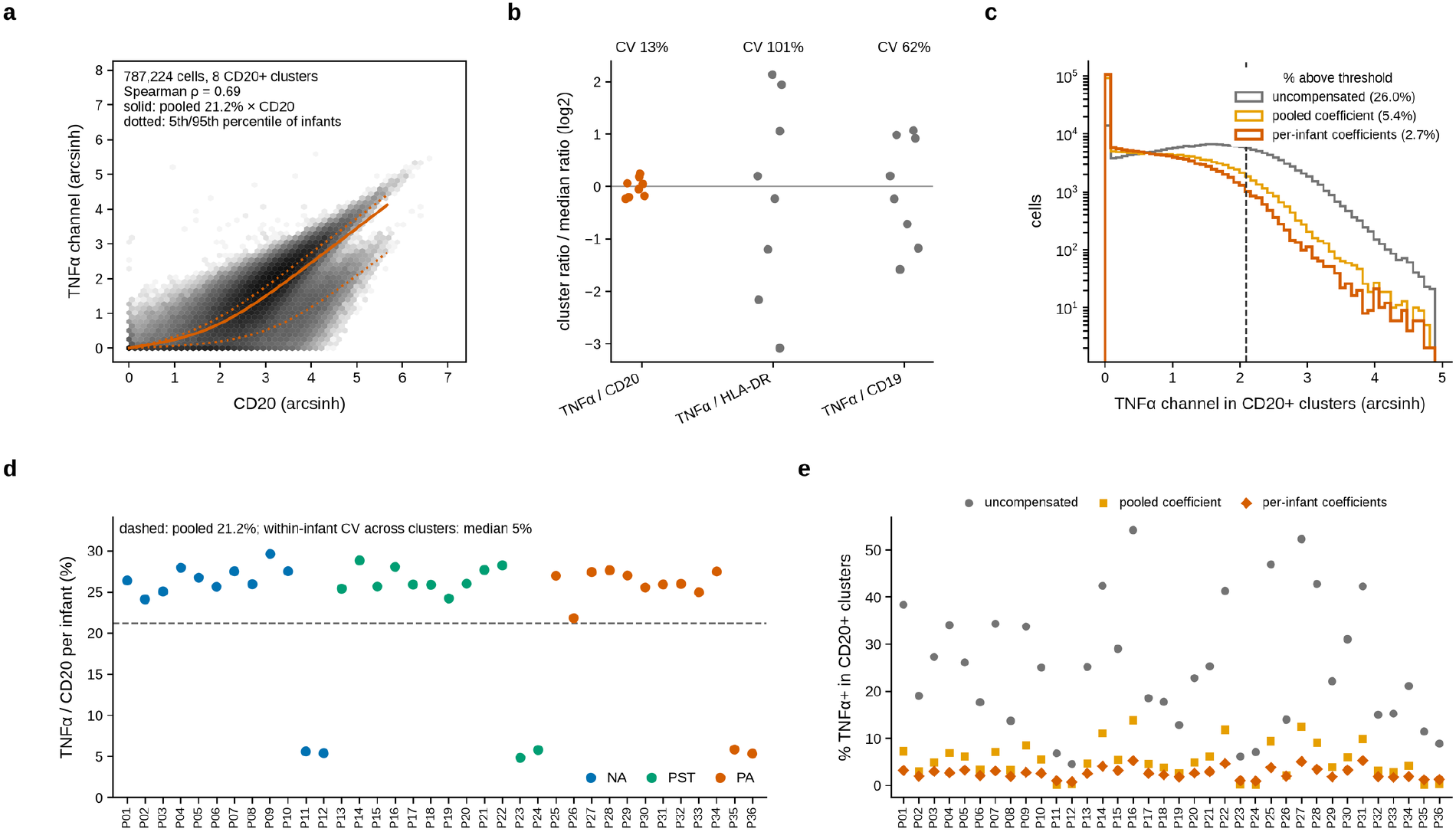
Spillover from CD20 into the TNFα channel. **a**, TNFα (^152^Sm) against CD20 (^147^Sm) for single cells of the eight CD20^+^ clusters (787,224 cells, 200,000 shown; arcsinh scale). The solid line marks the pooled spillover coefficient (21%); dotted lines mark the 5th and 95th percentiles of the per-infant coefficients. **b**, Ratio of TNFα to CD20, HLA-DR and CD19 in cluster medians of the same clusters, each relative to its median ratio; coefficients of variation are given above. **c**, TNFα channel in the same cells without compensation, compensated with the pooled coefficient and compensated with per-infant coefficients; dashed line, positivity threshold (20 counts). **d**, Spillover coefficient (fraction of the CD20 signal in the TNFα channel) per infant, median over its CD20^+^ clusters, colored by clinical group; dashed line, pooled coefficient. **e**, Percentage of TNFα^+^ cells in each infant’s CD20^+^ clusters without compensation, compensated with the pooled coefficient and compensated with the infant’s own coefficient.

**Extended Data Fig. 4.**
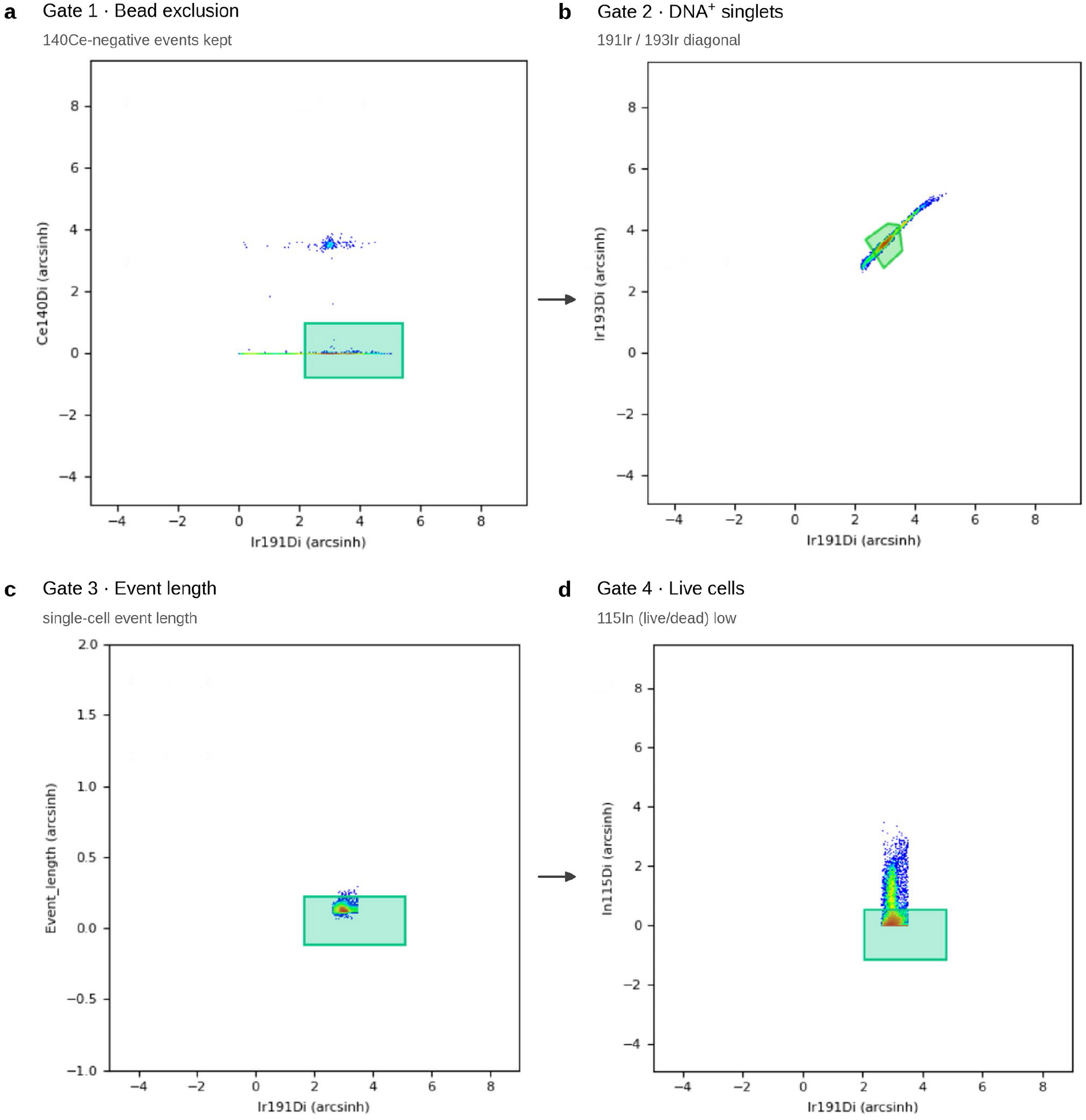
Gating strategy applied before clustering. After PeacoQC quality control, events from all 36 files were gated on cytowhiz.ai: **a**, exclusion of normalization beads (^140^Ce-positive) against the ^191^Ir DNA channel; **b**, DNA-positive single cells on ^191^Ir versus ^193^Ir; **c**, event length; **d**, live cells (^115^In DOTA-maleimide-negative). Plots show pooled events with axes as displayed by the platform (arcsinh scale); shaded regions are the gates.

**Extended Data Fig. 5.**
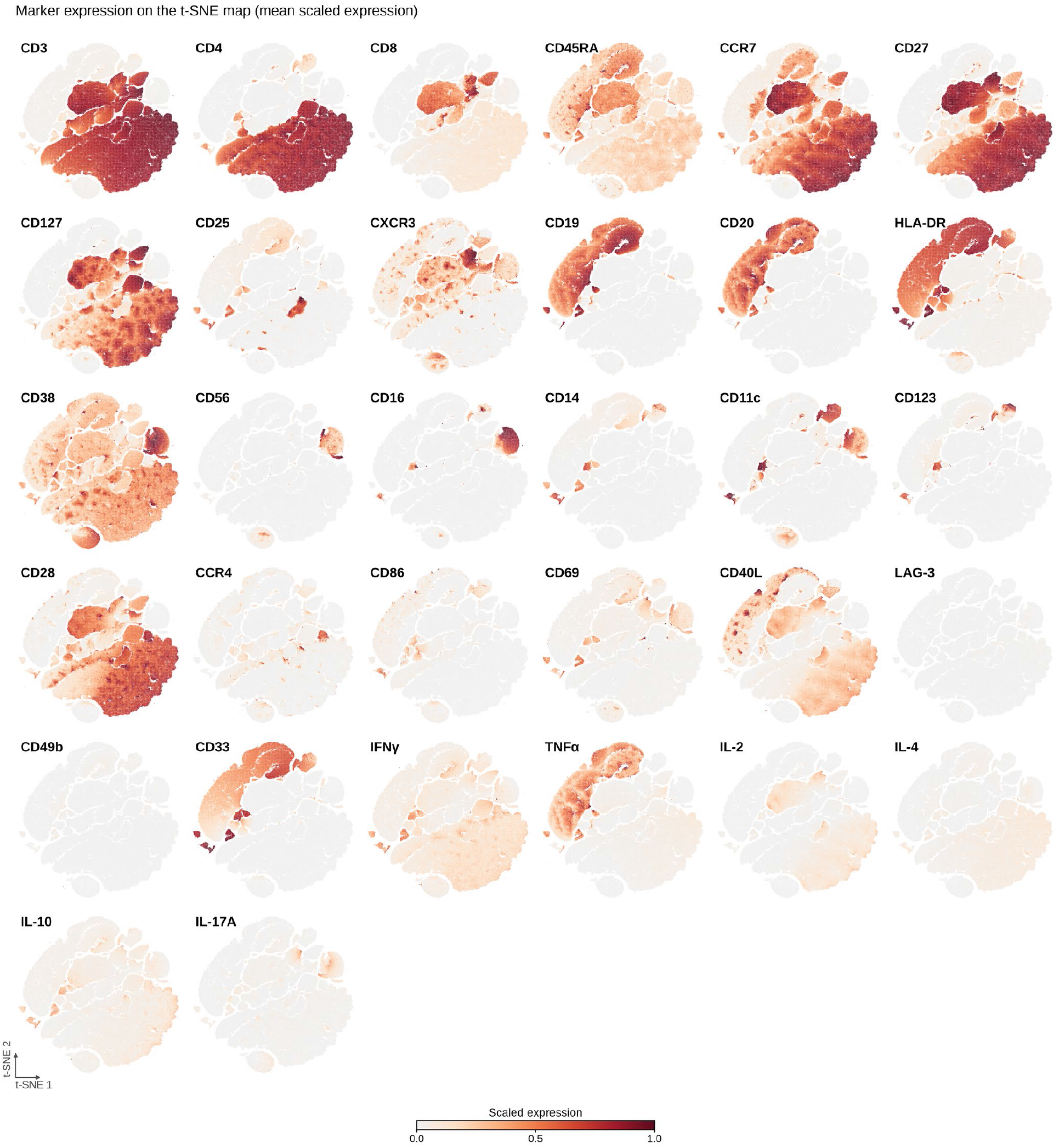
Expression of all 32 antibody channels on the t-SNE map. Mean scaled expression, computed from all cells; scaling as in Fig. 1d.

**Supplementary Table 1.**
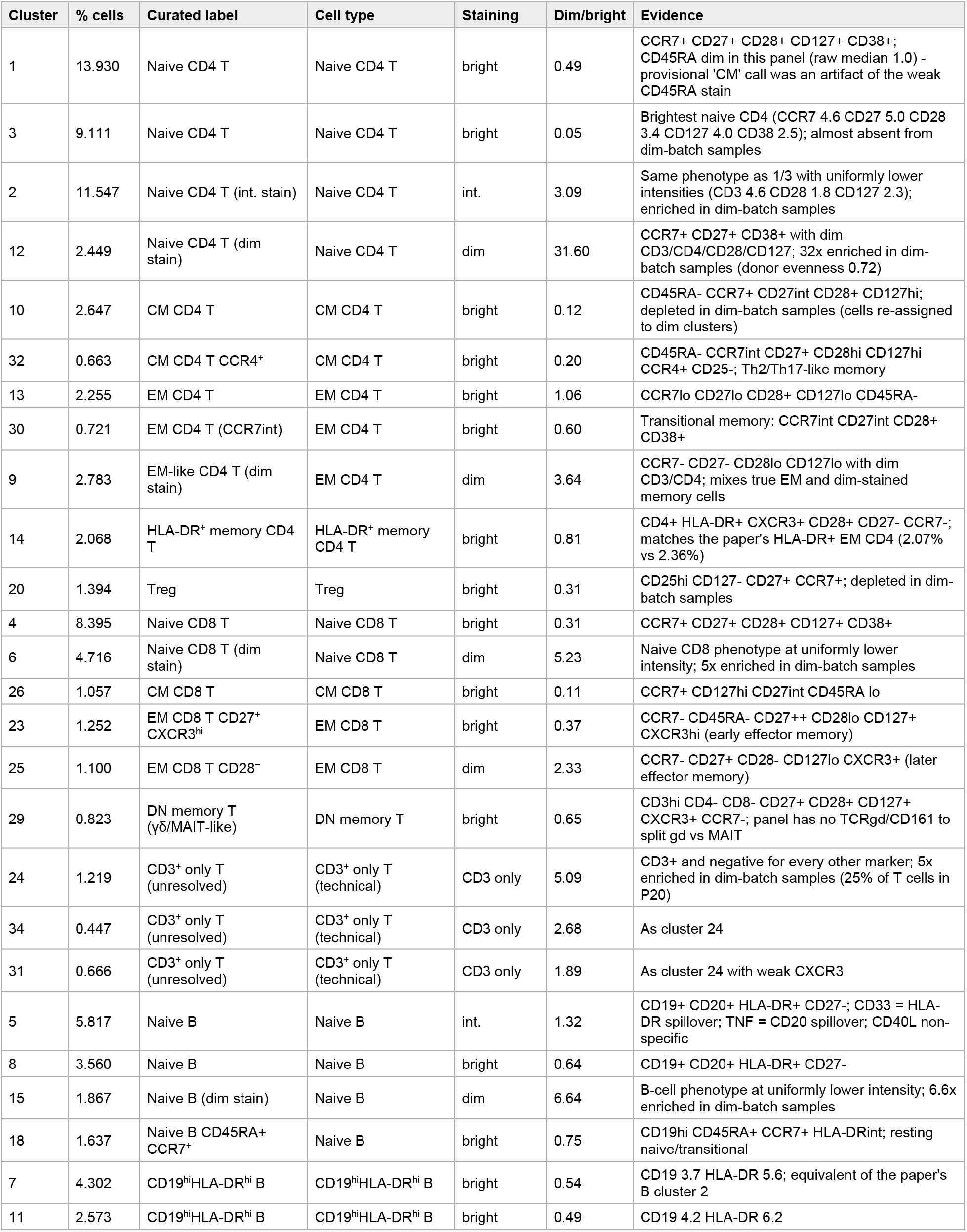
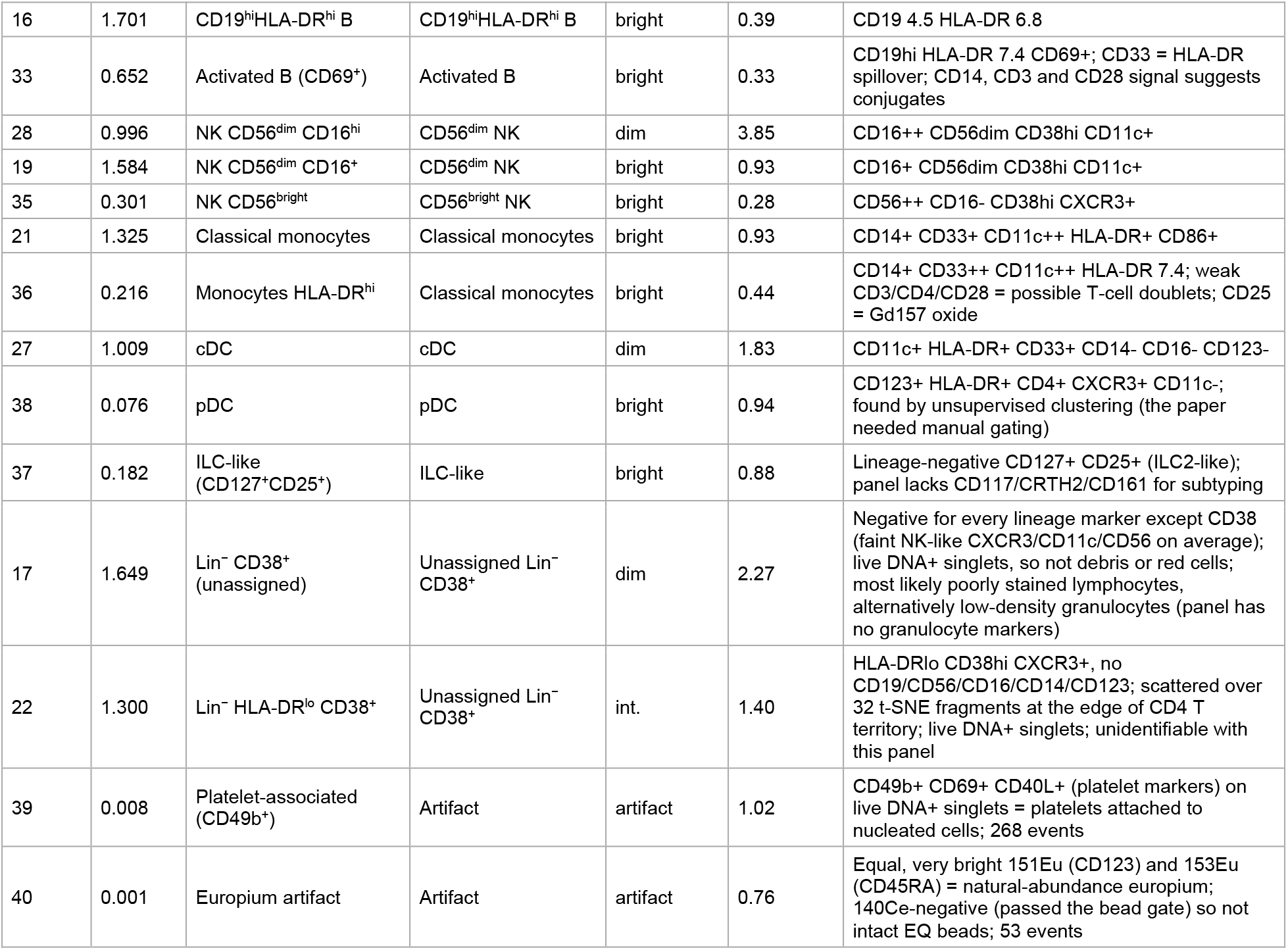
Curated annotation of the 40 PhenoGraph clusters. % cells, share of all 3,560,317 cells; staining, staining state relative to batch; dim/bright, ratio of the cluster’s mean frequency in dim-versus bright-batch samples.

**Supplementary Table 2.** Abundance statistics. Median % of cells per group; overall *q*, Benjamini–Hochberg-adjusted F-test of group in the batch-adjusted model; pairwise entries give the batch-adjusted fold change with its *q* value (Benjamini–Hochberg across all pairwise comparisons of the family); batch, frequency ratio dim/bright.

| Cell type | NA | PST | PA | Overall $q$ | PST vs NA | PA vs NA | PA vs PST | Batch |
| --- | --- | --- | --- | --- | --- | --- | --- | --- |
| Naive CD4 T | 38.80 | 31.57 | 39.88 | 0.17 | 0.76 (0.15) | 0.94 (0.71) | 1.23 (0.27) | 0.94 |
| CM CD4 T | 3.89 | 1.47 | 2.42 | 0.49 | 0.81 (0.53) | 0.75 (0.41) | 0.93 (0.82) | 0.14 |
| EM CD4 T | 5.28 | 6.80 | 4.71 | 0.60 | 1.09 (0.71) | 0.91 (0.68) | 0.83 (0.50) | 1.67 |
| HLA-DR<sup>+</sup> memory CD4 T | 1.75 | 2.48 | 1.61 | 0.045 | 1.45 (0.27) | 0.73 (0.35) | 0.51 (0.033) | 0.60 |
| Treg | 1.17 | 0.95 | 1.64 | 0.49 | 1.21 (0.50) | 1.25 (0.41) | 1.03 (0.91) | 0.30 |
| Naive CD8 T | 13.10 | 11.03 | 13.52 | 0.31 | 0.96 (0.80) | 1.17 (0.35) | 1.22 (0.27) | 1.08 |
| CM CD8 T | 1.16 | 0.31 | 0.63 | 0.62 | 0.61 (0.50) | 0.76 (0.69) | 1.25 (0.73) | 0.10 |
| EM CD8 T | 2.37 | 2.59 | 1.80 | 0.68 | 1.18 (0.66) | 0.96 (0.91) | 0.82 (0.59) | 0.90 |
| DN memory T | 0.83 | 1.13 | 0.79 | 0.31 | 1.83 (0.27) | 1.31 (0.56) | 0.72 (0.50) | 0.44 |
| Naive B | 10.23 | 13.73 | 13.34 | 0.34 | 1.22 (0.36) | 1.23 (0.35) | 1.01 (0.98) | 1.30 |
| CD19<sup>hi</sup>HLA-DR<sup>hi</sup> B | 8.19 | 5.45 | 9.93 | 0.49 | 1.05 (0.83) | 1.27 (0.40) | 1.21 (0.50) | 0.48 |
| Activated B | 0.67 | 0.44 | 0.82 | 0.62 | 1.19 (0.67) | 1.31 (0.50) | 1.10 (0.82) | 0.27 |
| CD56<sup>dim</sup> NK | 1.96 | 2.28 | 1.73 | 0.68 | 1.24 (0.65) | 1.24 (0.65) | 1.00 (1.00) | 1.31 |
| CD56<sup>bright</sup> NK | 0.23 | 0.13 | 0.28 | 0.49 | 1.19 (0.71) | 1.57 (0.41) | 1.32 (0.59) | 0.26 |
| Classical monocytes | 1.43 | 1.66 | 0.92 | 0.34 | 1.39 (0.50) | 0.78 (0.59) | 0.56 (0.27) | 0.62 |
| cDC | 0.57 | 0.97 | 0.42 | 0.086 | 1.69 (0.35) | 0.66 (0.41) | 0.39 (0.069) | 1.33 |
| pDC | 0.05 | 0.09 | 0.06 | 0.17 | 1.73 (0.15) | 1.29 (0.50) | 0.74 (0.41) | 0.90 |
| ILC-like | 0.15 | 0.19 | 0.21 | 0.17 | 1.43 (0.26) | 1.46 (0.22) | 1.02 (0.96) | 0.77 |
| Unassigned Lin<sup>-</sup> CD38<sup>+</sup> | 2.52 | 4.49 | 1.50 | 0.009 | 1.66 (0.14) | 0.73 (0.35) | 0.44 (0.006) | 1.42 |
| T cells (all) (total) | 72.82 | 68.36 | 71.77 | 0.24 | 0.91 (0.25) | 0.95 (0.45) | 1.05 (0.45) | 1.00 |
| CD4 T (total) | 54.22 | 45.42 | 50.52 | 0.24 | 0.86 (0.25) | 0.92 (0.45) | 1.07 (0.45) | 0.94 |
| CD8 T (total) | 16.51 | 14.63 | 17.43 | 0.46 | 0.97 (0.82) | 1.11 (0.45) | 1.14 (0.45) | 0.98 |
| B cells (total) | 19.33 | 20.60 | 22.98 | 0.24 | 1.13 (0.45) | 1.25 (0.25) | 1.10 (0.46) | 0.89 |
| NK cells (total) | 2.34 | 2.45 | 2.16 | 0.59 | 1.24 (0.48) | 1.30 (0.45) | 1.05 (0.87) | 1.13 |
| Monocytes (total) | 1.43 | 1.66 | 0.92 | 0.24 | 1.39 (0.45) | 0.78 (0.46) | 0.56 (0.26) | 0.62 |
| Dendritic cells (total) | 0.62 | 1.01 | 0.46 | 0.10 | 1.67 (0.28) | 0.69 (0.45) | 0.41 (0.083) | 1.29 |

